# Whole genome identification of potential G-quadruplexes and analysis of the G-quadruplex binding domain for SARS-CoV-2

**DOI:** 10.1101/2020.06.05.135749

**Authors:** Rongxin Zhang, Xiao Ke, Yu Gu, Hongde Liu, Xiao Sun

## Abstract

The Coronavirus Disease 2019 (COVID-19) pandemic caused by SARS-CoV-2 (Severe Acute Respiratory Syndrome Coronavirus 2) quickly become a global public health emergency. G-quadruplex, one of the non-canonical secondary structures, has shown potential antiviral values. However, little is known about G-quadruplexes on the emerging SARS-CoV-2. Herein, we characterized the potential G-quadruplexes both in the positive and negative-sense viral stands. The identified potential G-quadruplexes exhibits similar features to the G-quadruplexes detected in the human transcriptome. Within some bat and pangolin related beta coronaviruses, the G-quartets rather than the loops are under heightened selective constraints. We also found that the SUD-like sequence is retained in the SARS-CoV-2 genome, while some other coronaviruses that can infect humans are depleted. Further analysis revealed that the SARS-CoV-2 SUD-like sequence is almost conserved among 16,466 SARS-CoV-2 samples. And the SARS-CoV-2 SUD_core_-like dimer displayed similar electrostatic potential pattern to the SUD dimer. Considering the potential value of G-quadruplexes to serve as targets in antiviral strategy, we hope our fundamental research could provide new insights for the SARS-CoV-2 drug discovery.

## Introduction

The COVID-19 pandemic, which first broke out in China, has rapidly become a global public health emergency within a few months[1]. According to the statistics from the Johns Hopkins Coronavirus Resource Center (https://coronavirus.jhu.edu/map.html), 6.6 million cases have been confirmed, with a death toll rising to 389,000. Since 2000, humans have suffered at least three coronavirus outbreaks, and they were Severe Acute Respiratory Syndrome (SARS) in 2003[2-5], Middle East Respiratory Syndrome (MERS) in 2012[2, 5], and COVID-19. Scientists identified and sequenced the virus early in this outbreak, and named it SARS-CoV-2[6]. The symptoms of the patients infected with the novel coronavirus vary from person to person, and fever, cough, and fatigue are the most common ones[7-10]. The clinical chest CT (Computed tomography) and nucleic acid testing are the most typical methods of diagnosing COVID-19[9, 10]. It is worth noting that the recent achievements in AI (Artificial Intelligence) aid diagnosis technology[11] and CRISPR-Cas12-based detection methods[12] are expected to expand the diagnosis of COVID-19. Despite the great efforts of the researchers, so far, no specific clinical drugs or vaccines have been developed to cope with COVID-19.

SARS-CoV-2 is a Betacoronavirus within the Coronaviridae family that is the culprit responsible for the COVID-19 pandemic[13, 14] (Fig. 1A). Studies have confirmed that SARS-CoV-2 is a positive-sense single-stranded RNA ((+)ssRNA) virus with a total length of approximately 30k. The positive-sense RNA strand of SARS-CoV-2 can serve as a template to produce viral proteins related to replication, structure composition, and other functions or events[15, 16]. One of the hotspots is how the SARS-CoV-2 entry the host cells. SARS-CoV-2 has shown a great affinity to the angiotensin-converting enzyme 2 (ACE2), which has been proved to be the binding receptor for SARS-CoV-2[17, 18]. After entering the host cells, the viral genomic RNA will be released to the cytoplasm, and the ORF1a/ORF1ab are subsequently translated into replicase polyproteins of pp1a/pp1ab, which will be cleaved into some non-structural proteins (nsps). These non-structure proteins ultimately form the replicase-transcriptase complex for replication and transcription. Along with the full-length positive and negative-sense RNAs, a nested set of sub-genomic RNAs (sgRNAs) are also synthesized, and mainly translated into some structural proteins and accessory proteins. When assembly finished, the mature SARS-CoV-2 particles are released from infected host cells via exocytosis[19]. Mounting evidence suggests that bats and pangolins are the suspected natural host and intermediate host of SARS-CoV-2[20-23]. Intriguingly, a report from Yongyi Shen et al. showed that SARS-CoV-2 might be the recombination product of Bat-CoV-RaTG13-like virus and Pangolin-CoV-like virus[24].

**Fig. 1.**
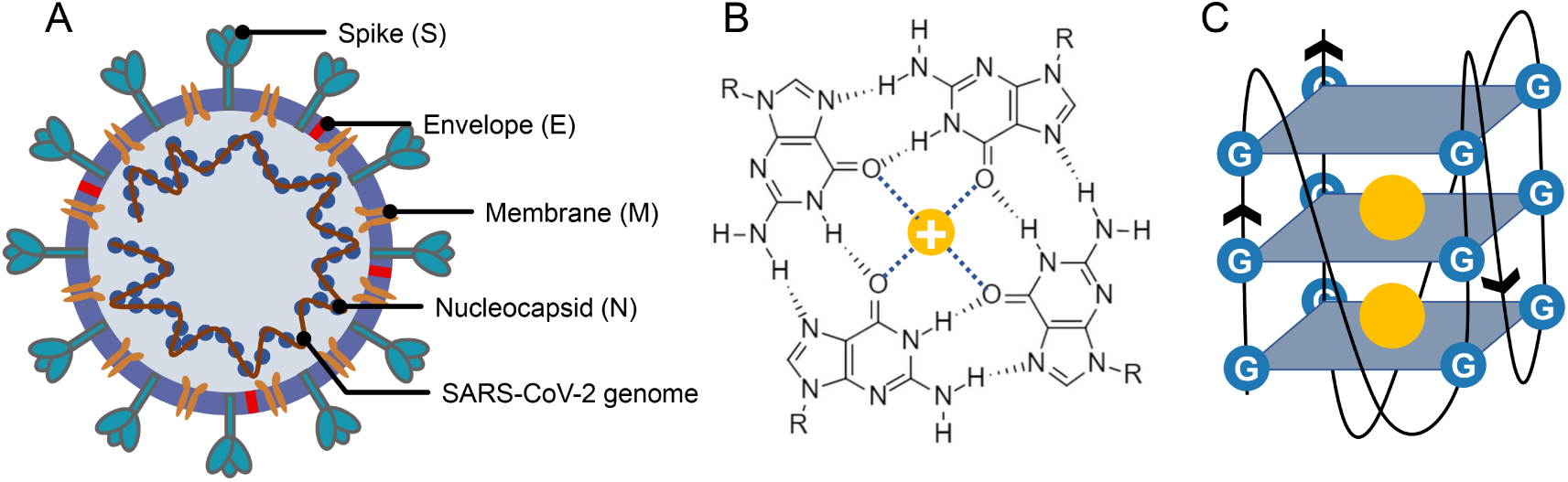
Structure of SARS-CoV-2, G-quartet, and G-quadruplex. (A) The SARS-CoV-2 particle structure is composed of four structural proteins, which are the spike protein, the envelope protein, the membrane protein, and the nucleocapsid protein. The nucleocapsid proteins are bound to the SARS-CoV-2 genome. (B) Structure of G-quartet, the neighboring guanines are connected via Hoogsteen hydrogen. The cation is indicated by a yellow circle. (C) The G-quadruplex is formed by stacking multiple G-quartets, and the stabilization of the structure is partially determined by the central cations.

G-quadruplexes are the non-canonical nucleic acid structures usually formed in G-rich regions both in DNA and RNA strands[25-27]. The G-quadruplex is formed by stacking G-quartets (Fig. 1B) on top of each other, in which the four guanines making up a G-quartets are connected via Hoogsteen pairs (Fig. 1C)[25-28]. Extensive research indicated that G-quadruplexes were involved in many critical biological processes, including DNA replication[29-32], telomere regulation[33-37], and RNA translation[38-41]. It has been proved that G-quadruplexes existed in the viral genome and can regulate the viral biological processes, which made it possible to function as potential drug targets for antiviral strategy[42-44]. A study made by Jinzhi Tan et al. demonstrated that the SARS-Unique Domain (SUD) within the nsp3 (non-structural protein 3) of SARS coronavirus (SARS-CoV) exhibits the binding preference to the G-quadruplex structure in the human transcript, and potentially interfere with host cell antiviral response[45]. They also identified several amino acid residues that were tightly associated with its binding capacity. Yet, whether the SARS-CoV-2 contains a SUD-like structure and whether the structure sequence is conserved among SARS-CoV-2 samples needs further interpretation. Besides, the G-quadruplexes in some well-known virus, such as HIV-1[46-49] (Human Immunodeficiency Viruses type 1), ZIKV[50] (ZIKA Virus), HPV[51, 52] (Human Papillomavirus) and EBOV[53] (Ebola virus) have been studied. However, our understanding of the G-quadruplexes, their potential roles, and the SUD-like structures in the emerging SARS-CoV-2 are lacking.

In this study, we depicted the potential G-quadruplexes (PG4s) in the SARS-CoV-2 by combining several G-quadruplex prediction tools. The PG4s in SARS-CoV-2 presented similar features to the two-quartet G-quadruplexes in the human transcriptome, which potentially supported the formation and existence of the G-quadruplexes in SARS-CoV-2. Additionally, we investigated the difference in selective constraints between the G-quartets and other nucleotides in the SARS-CoV-2 genome. To further elucidated the possible pathogenic mechanism of SARS-CoV-2, we examined the SUD-like sequence and structure in SARS-CoV-2 that are critical to binding the G-quadruplexes in host transcripts.

## Results

### Whole genome identification and annotation of potential G-quadruplexes

To get the potential G-quadruplexes in the SARS-CoV-2 genome, we took the strategy described as follows (Fig. 2A): (i) Predicting the PG4s with three software independently. (ii) Merging the prediction results of the PG4s and evaluating the G-quadruplex folding capabilities by the cG/cC scores. (iii) The PG4s with cG/cC scores higher than the threshold were selected as candidates for further analysis. Here, the threshold for determining whether PG4s can be folded was set to 2.05, as described in the study of Jean-Denis Beaudoin et al.[54] In total, we obtained 24 PG4s (Table. 1) in the positive or negative-sense strands for further analysis.

**Table. 1.**
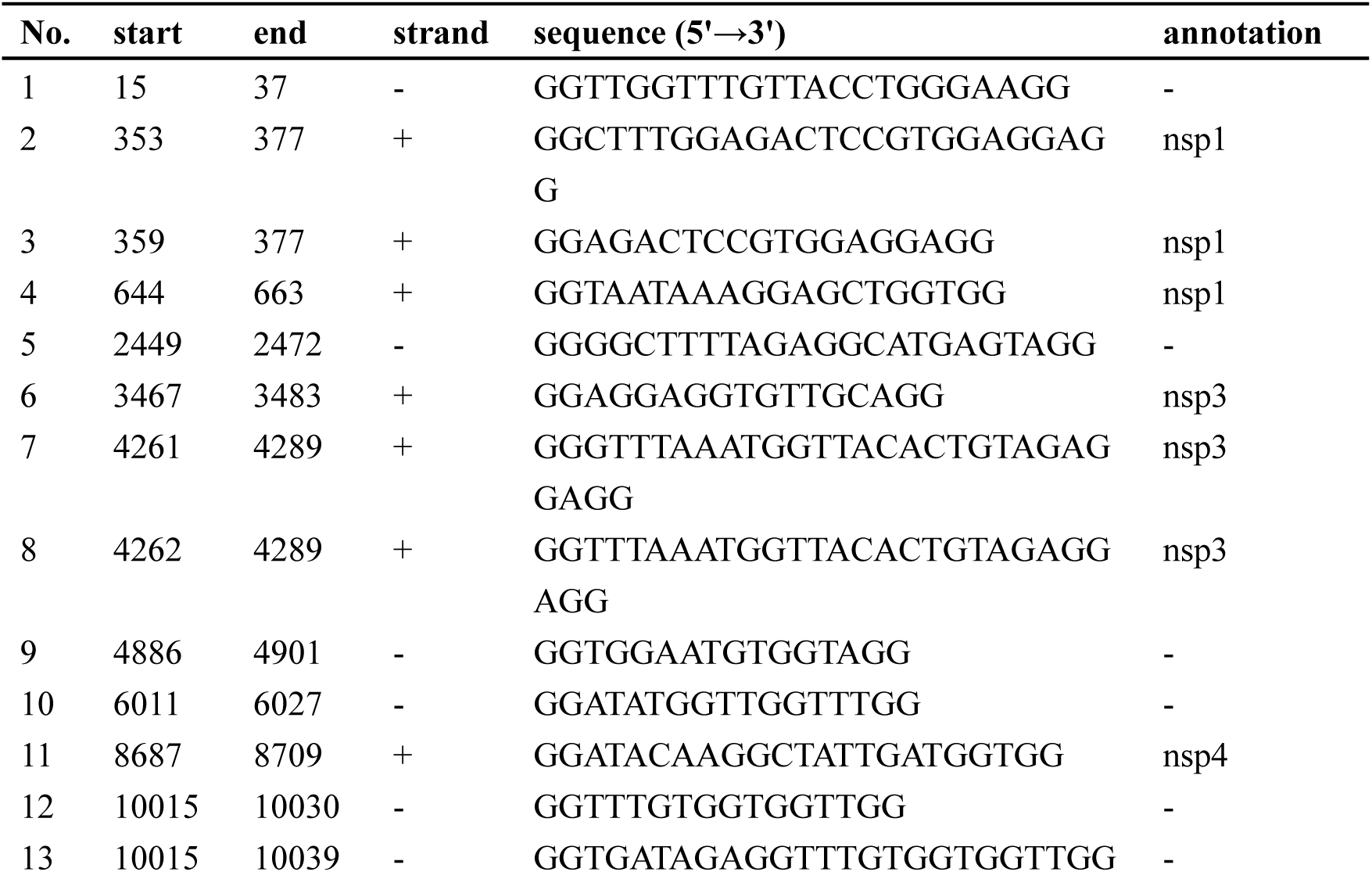
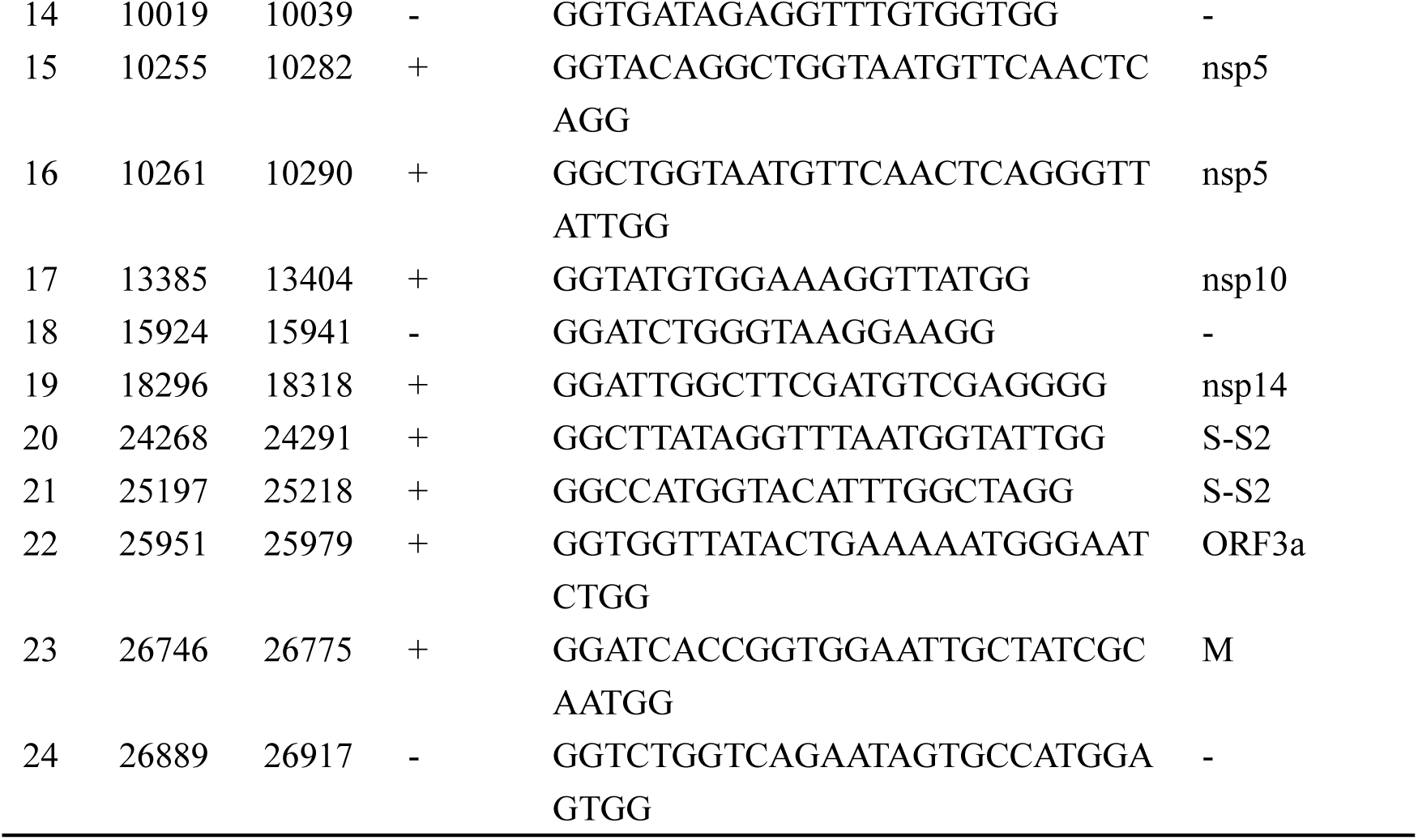
The PG4s found in the SARS-CoV-2.

**Fig. 2.**
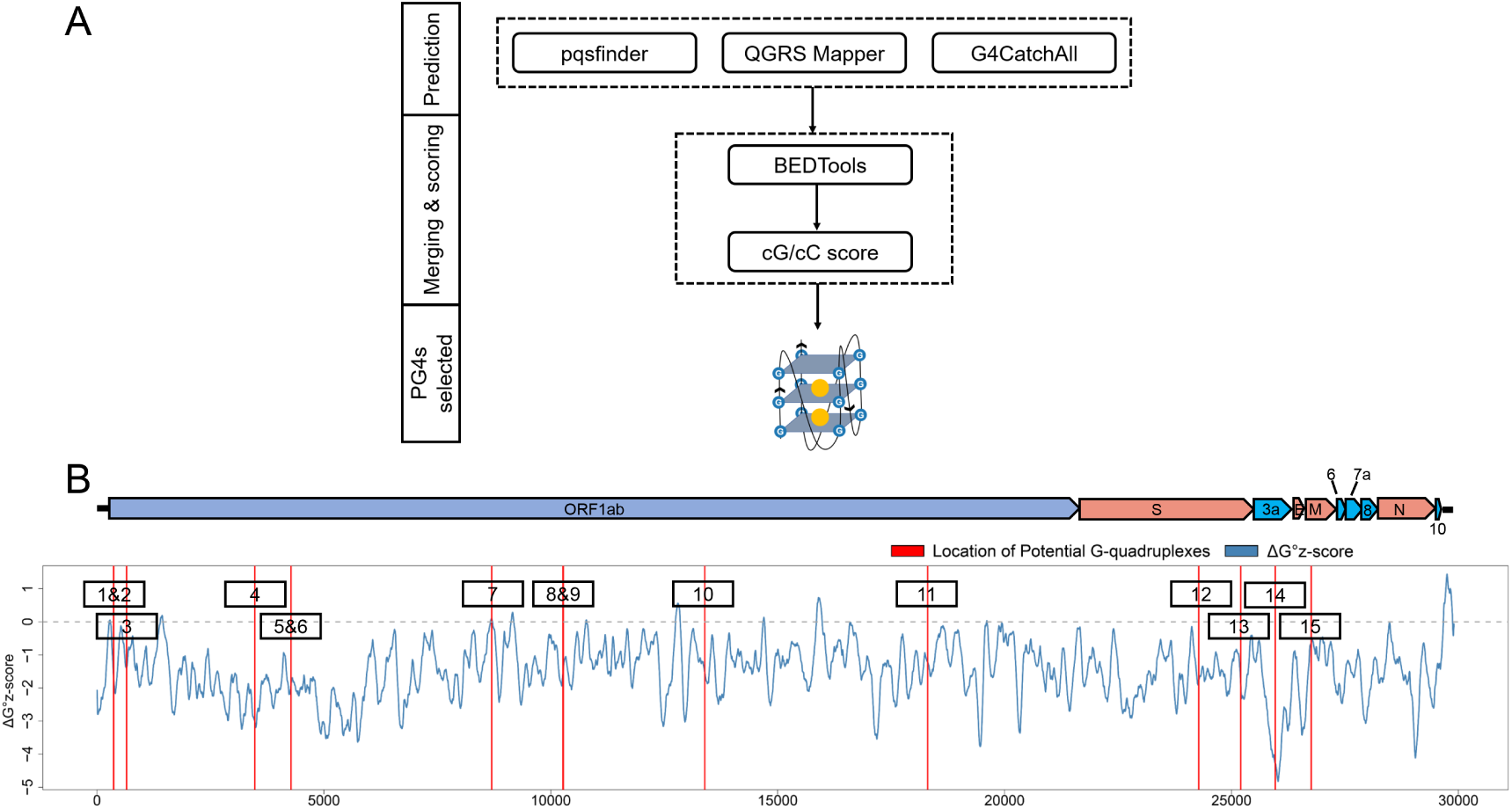
Detection and annotation of the PG4s. (A) The schematic flow of PG4s detection. The G-quadruplex prediction tools, pqsfinder, QGRS Mapper, and G4CatchAll were utilized for the prediction of PG4s. BEDTools and the cG/cC scoring system were applied to merging and scoring the PG4s. After screening, the PG4s that used in this study were generated. (B) Visualization of the PG4s in the SARS-CoV-2 genome. Top panel, the genome organization of SARS-CoV-2. Bottom panel, the average Δ G° z-score for each nucleotide (blue curve) in the SARS-CoV-2 genome, the location of PG4s are plotted with red vertical lines. The order of the PG4s is marked with a black box. Please note that only the PG4s in the positive-sense strand are visualized.

To annotate the PG4s, the reference annotation data (in gff3 format) of SARD-CoV-2 were downloaded from the NCBI database with the accession number of NC_045512. Firstly, we focused on the PG4s on the positive-sense strand. Fifteen of the 24 PG4s (67.5%) were located on the positively-sense strand, the vast majority of them were harbored in non-structural proteins including nsp1, nsp3, nsp4, nsp5, nsp10 and nsp14, with the remaining ones located in the spike protein, orf3a, and the membrane protein. Secondly, we examined the PG4s on the negative-sense strand, which is an intermediate product of replication. Nine PG4s were scattered on the negative-sense strand.

To further characterize the potential canonical secondary structures competitive with G-quadruplexes, the landscape of thermodynamic stability of the SARS-CoV-2 genome was depicted by using ΔG° z-score[55]. In general, a positive ΔG° z-score implies that the secondary structure of this region tends to be less stable than the randomly shuffled sequence with the identical nucleotide composition, while a negative ΔG° z-score signifies higher stability than the randomly shuffled sequence. For each nucleotide in the SARS-CoV-2 genome, the ΔG° z-score was calculated for all the 120 nt windows covering the nucleotide, and an average ΔG° z-score was deduced then. Several PG4s are located in positions with a locally higher average ΔG° z-scores (Fig. 2B) which implied the relative instability of a canonical secondary structure and the lower possibility to adopt such a competitive structure against the G-quadruplex, which may ultimately favor the formation of G-quadruplex.

### Potential G-quadruplexes in SARS-CoV-2 show analogical features with the rG4s in the human transcriptome

In 2016, Chun Kit Kwok and co-workers profiled the RNA G-quadruplexes in the HeLa transcriptome by using the RNA G-quadruplex sequencing (rG4-seq) technology, and quantified the diversity of these RNA G-quadruplexes[56]. We set out to address the question of whether the potential G-quadruplexes in SARS-CoV-2 showed analogical features with the G-quadruplexes found in the human transcriptome and if these PG4s have the ability to form G-quadruplex structures. We noticed that the PG4s in SARS-CoV-2 are all in the two-quartet style. Therefore we retrieved the two-quartet RNA G-quadruplex sequence data generated in the rG4-seq experiment under the condition of K^+^ and pyrdiostatin (PDS). However, for some RTS (Reverse Transcriptase Stalling) sites labeled as two-quartet, there may exist overlapping G-quadruplexes with different loops (e.g., <u>GG</u>CACAGCA<u>GG</u>CATC<u>GG</u>A<u>GG</u>TGA<u>GG</u>C<u>GGGG</u>), and it is difficult to determine which one was formed in the experiment. In order to eliminate the ambiguity, only the RTS sites containing non-overlapping two-quartet G-quadruplex (e.g., GTCATTTTTTGTGTTT<u>GG</u>TTT<u>GG</u>T<u>GG</u>T<u>GG</u>C) were considered.

Firstly, we investigated the loop length distribution pattern of the two-quartet PG4s in both SARS-CoV-2 and the human transcriptome (Fig. 3A). As a whole, the two-quartet PG4s in SARS-CoV-2 and the human transcriptome displayed similar loop length distribution patterns, and the loop length of the PG4s in SARS-CoV-2 falls into the scope of the ones from the human transcriptome. The distributions of loop length between the SARS-CoV-2 PG4s and the human two-quartet G-quadruplexes did not show discrepancies (Fig. S1, Wilcoxon test, *p*-value = 0.4552).

**Fig. 3.**
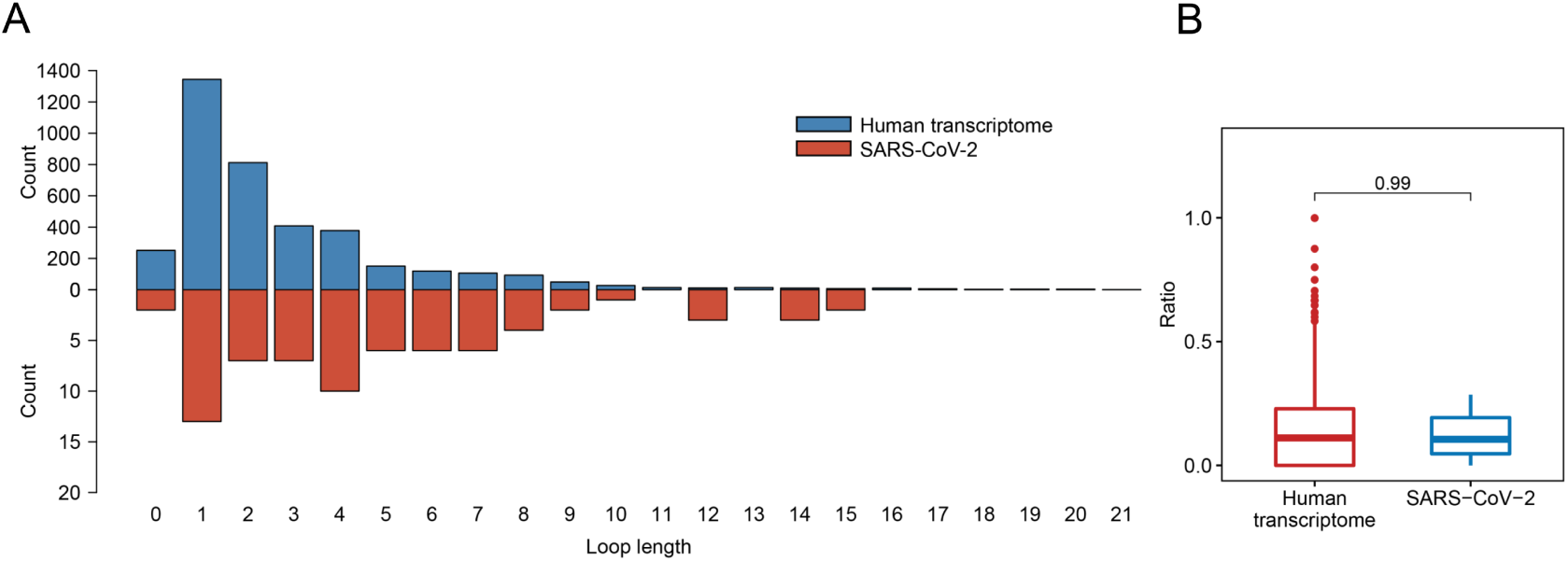
Feature comparison of potential G-quadruplexes found in rG4-seq and SARS-CoV-2. (A) The Histogram represents the number of two-quartet G-quadruplex loops with different lengths in the human transcriptome and SARS-CoV-2, respectively. (B) The red boxplot shows the ratio of cytosine in human transcriptome two-quartet G-quadruplex loops. And the blue boxplot displays the ratio of cytosine in SARS-CoV-2 PG4 loops.

Considering the fact that the presence of multiple-cytosine tracks may hinder the formation of G-quadruplexes[54, 57], we examined the cytosine ratio in G-quadruplex loops (Fig. 3B). No significant difference in loop cytosine ratios was observed between the SARS-CoV-2 PG4s and the human two-quartet G-quadruplexes (Wilcoxon test, *p*-value = 0.9911), which suggested that the loop cytosine ratios between the two types of G-quadruplex were similar.

Taken together, our results suggested that the PG4s in SARS-CoV-2 displayed similar features to the rG4s in the human transcriptome.

### Potential G-quadruplexes are under selective constraints in bat and pangolin related beta coronaviruses

Recent research revealed that the G-quadruplexes in human UTRs (Untranslated Regions) are under selective pressures[58], and some coronaviruses on bats and pangolins are closely related to SARS-CoV-2. The conservation of the potential G-quadruplexes in the SARS-CoV-2 genome under selective constraints were analyzed. We collected some beta coronavirus genomic sequences of bats and pangolins from several public databases and used the NJ (Neighbor-Joining) method to construct the phylogenetic tree with 1,000 bootstrap replications (Fig. S2). The RS (Rejected Substitutions) score for each site in the SARS-CoV-2 reference genome was evaluated by using the GERP++ software.

We checked the RS score difference between the G-tract (continuous runs of G) nucleotides and other nucleotides. A significant discrepancy was observed, which means that the G-tracts nucleotides exhibit heightened selective constraints than other nucleotides in the SARS-CoV-2 genome (Fig. 4A, Wilcoxon test, *p*-value = 9.254 × 10^−8^). Considering that the G-tracts are composed of guanines, the conservation of guanines in and outside the G-tracts in the SARS-CoV-2 genome were also compared. We found that the guanines in G-tracts are under heightened selective constraints (Fig. 4B, Wilcoxon test, *p*-value = 3.363×10^−3^). The nucleotides within G-tracts are more relevant to the G-quadruplexes structural maintenance than loops. Then we compared the G-tract and loop RS scores. As expected, the G-tract RS scores were significantly higher than loops (Fig. 4C, Wilcoxon test, *p*-value = 3.962 × 10^−7^), which suggests that the G-tracts experienced stronger selective constraints.

**Fig. 4.**
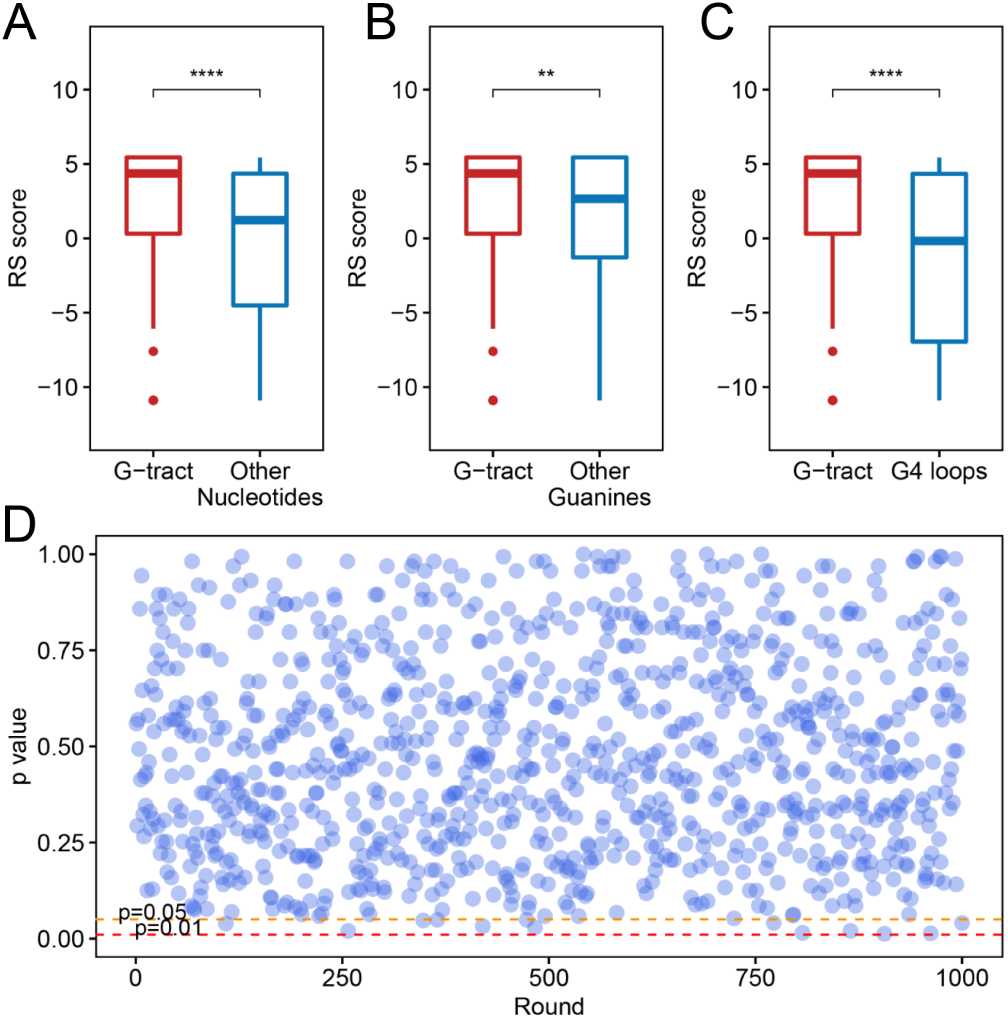
Potential G-quadruplexes exhibit heightened selective constraints in bat and pangolin related coronavirus. (A-C) Boxplot showing the difference of nucleotide RS scores in G-tract, other nucleotides, other guanines, and PG4 loops (\*\**p* ≤ 0.01, \*\*\*\**p* ≤ 0.0001). (D) Tests of the RS score difference between the fragments containing PG4s and randomly selected fragments. The abscissa indicates the round of the test, while the ordinate represents the *p*-value for each round.

We also checked that if the PG4s that are under heightened selective constraints is relevant to its inherent properties or potential functions rather than the sequence contexts. A random test was performed to check whether the fragments containing PG4s manifested different average RS scores compared with random fragments in the SARS-CoV-2 genome. The fragments containing PG4 were designated as the sequence 100 nt upstream and downstream of the PG4 centers. We conducted 1,000 rounds of tests. In each test, we randomly selected 50 fragments from the SARS-CoV-2 genome with a length of 200 nt and carried out the Wilcoxon test to assess the average RS score difference among the randomly selected fragments and the fragments containing PG4s. The *p*-value for each round was retained. As a result, no evident difference was observed as few *p*-values (13/1000) were less than 0.05 (Fig. 4D), suggesting that PG4s that are under heightened selective constraints is more likely to be related to its inherent properties or potential functions rather than sequence contexts.

### SARS-CoV-2 contains similar SUD to SARS-CoV

Both SARS-CoV and SARS-CoV-2 could cause acute disease symptoms, and the above coronavirus shares similar nucleic acid sequence compositions. There is a SUD in the SARS-CoV genome that can binding to the G-quadruplex structures and it is unclear if the SARS-CoV-2 genome possess the resemble structure. Thus, we started to explore whether the SARS-CoV-2 genome contains the protein-coding sequence similar to SUD and whether SARS-CoV-2 retains the ability to bind RNA G-quadruplexes. We collected the ORF1ab amino acid sequences of some coronaviruses, including seven known coronaviruses, which can infect humans and other coronaviruses belonging to different genera. Surprisingly, the SUD protein sequence is absent in some coronaviruses, especially in alpha, gamma, and delta coronaviruses (Fig. S3). In contrast, the SUD protein sequence is retained in several beta-coronavirus, particularly in bat and pangolin associated beta coronavirus. Moreover, among the seven coronaviruses that can infect humans, only SRAS-CoV and SARS-CoV-2 keep the SUD sequence, while the SUD sequence in MERS-CoV, HCoV-229E, HCoV-NL63, HCoV-OC43 and HCoV-HKU1 is depleted. Next, we examined eight key amino acid residues in SUD that previously reported associated with G-quadruplex binding affinity (Fig. 5A). Almost all the key amino acid residues are reserved in SARS-CoV-2, except one conservative replacement of K (Lysine) > R (Arginine). We hypothesized that if the G-quadruplex binding ability is essential for the SARS-CoV-2, the above amino acid residues should be conservative. We then investigated the conservation of the eight amino acid residues within SARS-CoV-2 samples. We retrieved the sequence alignment file of 16,466 SARS-CoV-2 samples from the GISAID database and calculated the mutation frequency for each nucleotide. We observed the frequency of nucleotide mutations in the above eight codons. As a result, a limited mutation frequency was found as compared to the whole genome average mutation frequency (Fig. 5B, frequency = 3.96). Although eight mutations were detected in glutamate (2432 E), seven of them were synonymous mutations. Next, we checked the electrostatic potential pattern in the SARS-CoV-2 SUD_core_-like dimer structure. The SARS-CoV-2 SUD_core_-like dimer structure is defined as the dimer structure formed by the amino acid residues in SARS-CoV-2 corresponding to the SUD of SARS. We found that the SUD_core_-like dimer of SARS-CoV-2 and the SUD_core_ of SARS present analogical electrostatic potential patterns. The positively charged patches were observed in the core of the SUD_core_-like dimer, which was surrounded by negatively charged patches (Fig. 5C). In contrast, when the dimer is rotated 180°, a slightly inclined narrow cleft with negative potential accompanied by the positively charged patches was discovered (Fig. 5D). And the above patterns also appeared in the SUD dimer. In the previous reports, several positively charged patches located in the center and back of the dimer were presumed to bind the G-quadruplex structures. By comparison with the electrostatic potential of the SARS SUD_core_ dimer, we identified the positively charged patches located in the center and back of the SARS-CoV-2 SUD_core_-like dimer, which can potentially bind the G-quadruplexes (Fig. 5C-D).

**Fig. 5.**
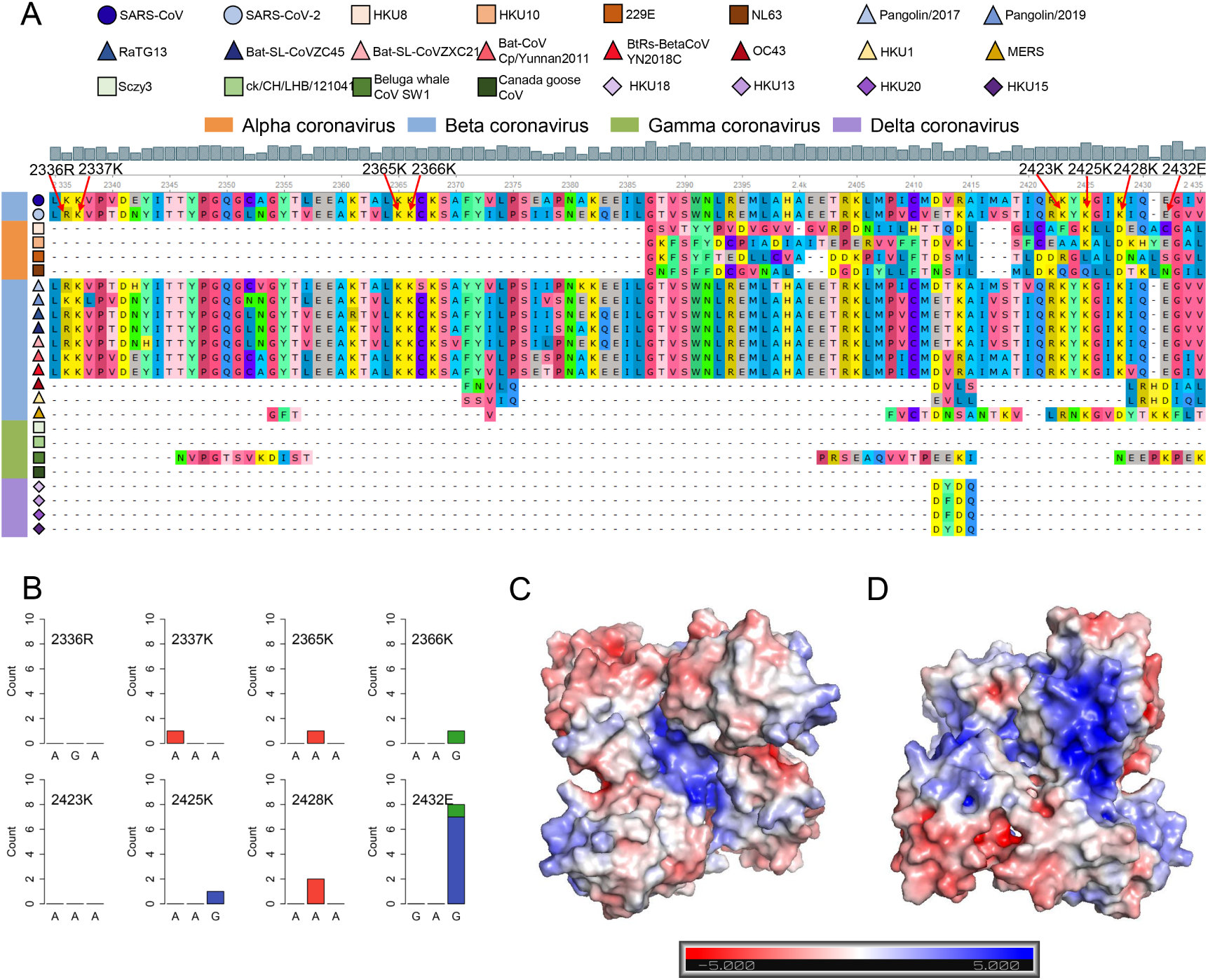
SARS-CoV-2 contains the SUD_core_-like sequence and dimer structure. (A) Sequence alignment of four different genera coronaviruses. The shapes in various colors mark different kinds of coronaviruses. The color bar represents different genera of coronaviruses (orange, alpha coronavirus; blue, beta coronavirus; green, gamma coronavirus; purple, delta coronavirus). The grey histogram shows the consensus of the alignment sites. The eight amino acid residues related to the G-quadruplex binding affinity are labeled by red arrows. (B) Nucleotide mutation count of eight amino acid residues among 16,466 SARS-CoV-2 samples. (C-D) The electrostatic potential surface of the SARS-CoV-2 SUDcore-like dimer with different orientations (the front (C) and rotate 180°(D)). The blue and red showed positive and negative potential, respectively.

## Discussion

The COVID-19 pandemic has caused huge losses to humans and made people pay more attention to public health. A large number of scientists all over the world have been engaged in the fight against the outbreak. The SARS-CoV-2 coronavirus is the key culprit responsible for the outbreak, and no specific inhibitor drugs have been developed yet. G-quadruplexes have shown tremendous potential for the development of anticancer[59-62] and antiviral drugs[44, 63, 64], as G-quadruplexes can interfere with many biological processes that are critical to cancer cells and viruses. Therefore, it is necessary to quantify and characterize the PG4s in the SARS-CoV-2 genome to provide a possible novel method for the treatment of COVID-19.

In this study, besides three popular G-quadruplexes prediction tools, the cG/cC scoring system, which is specially designed for the identification of RNA G-quadruplexes, was adopted to determine the PG4s. Indeed, we did not find the G-quadruplexes with three or more G-quartets, which are generally considered to be more stable than the two-quartet G-quadruplexes. One of the controversial issues lies on the stability of the two-quartet G-quadruplexes, especially the folding capability of those G-quadruplexes *in vivo*. However, it is well-acknowledged that the RNA G-quadruplexes is more stable than their DNA counterparts[65, 66] and SARS-CoV-2 is a single-strand RNA virus, which may be conducive to its structure formation. Several emerging studies have demonstrated the formation of two-quartet G-quadruplexes in viral sequences, which could serve as antiviral elements under the presence of G-quadruplex ligands[53, 67, 68]. Moreover, the K^+^ (potassium ion), one of the primary positive ions inside human cells, can strongly support the formation of G-quadruplexes. Nevertheless, whether the SARS-CoV-2 G-quadruplexes could form *in vivo* requires overwhelming proofs.

Most of the PG4s we detected were located in the positive-sense strand. The G-quadruplex forming sequences in the SARS-CoV genome were presumed to function as the chaperones of SUD, and their interaction was essential for the SARS-CoV genome replication[69]. ORF1ab that encodes the replicase proteins is required for the viral replication and transcription. Some PG4s were found to harbored in ORF1ab, and whether these PG4s were related to the replication of the viral genome and interact with SUD-like structures like in SARS-CoV, is worthy of further investigation. In addition to ORF1ab, there exists several PG4s in the structural and accessory protein-coding sequences as well as the sgRNAs that containing the above protein sequences. Some studies have characterized the impact of G-quadruplex structures on the translation of human transcripts, and an apparent inhibitory effect was observed [38, 57, 70]. The translation of some SARS-CoV-2 proteins requires the involvement of human ribosomes; thus, it is possible to repress the translation of SARS-CoV-2 proteins via stabilizing the G-quadruplex structures. In fact, this inhibition effect has been reported in some other viral studies [67, 71]. The negative-sense strand serves as templates for the synthesis of the positive-sense strand and the sub-genomic RNAs. The identified potential G-quadruplexes were broadly distributed in the negative-sense strand. Notably, we observed one PG4 located at the 3’ end of the negative-sense strand. A previous study confirmed that the stable G-quadruplex structures located at the 3’ end of the negative-sense strand could inhibit the RNA synthesis by reducing the activity of the RdRp (RNA-dependent RNA polymerase) [72]. Therefore, it is necessary to further investigate whether the PG4 at the 3’ end of the negative-sense strand of SARS-CoV-2 could inhibit RNA synthesis. In addition, recent research revealed that the high-frequency trinucleotide mutations (G28881A, G2882A and G28883C) were detected in the SARS-CoV-2 genome [73, 74]. G28881A and G28882A always co-occur within the same codon, which means a positive selection of amino acid [75]. We noticed that the trinucleotide mutations were in the G-rich sequence from 28881 nt to 28917 nt (5’ <u>GGGG</u>AACTTCTCCTGCTAGAAT**<u>GG</u>CT<u>GG</u>CAAT<u>GG</u>C<u>GG</u>** 3’). The potential G-quadruplex downstream of the trinucleotide mutations was filtered by the cG/cC score system as the presence of cytosine tracks within and flanking of the potential G-quadruplex reduce the cG/cC score; however, in fact, this potential G-quadruplex showed a relative lower MFE (Minimum Free Energy) among all the potential G-quadruplexes we detected. The consequence of the trinucleotide mutations was still elusive. Whether the mutations have an internal causality with the G-rich sequence still needs to be elucidated.

The SUD in SARS, which is thought to be related to its terrible pathogenicity, has displayed binding preference to the G-quadruplexes in human transcripts[45]. Our analysis revealed that the novel coronavirus SARS-CoV-2 contained a similar domain to SUD as well. Furthermore, several amino acid residues previously reported to be an indispensable part of the G-quadruplexes binding capability are retained in SARS-CoV-2. Further exploration indicated that the eight key amino acid residues were conserved in numerous SARS-CoV-2 samples across countries all over the world, suggesting the essentiality of the above residues. It is supposed that the binding of SUD to G-quadruplexes could affect transcripts stability and translation, hence impairing the immune response of host cells. The expression of host genes in SARS-CoV-2 infected cells is extremely inhibited[15]; therefore, we speculate that the SARS-CoV-2 may possess the similar mechanism with SARS-CoV that can inhibit the expression of some important immune-related genes to escape immune defense. Herein, we briefly depict the possible role of G-quadruplexes in the antiviral mechanism and pathogenicity, and the development of certain G-quadruplex specific ligands might be a promising antiviral strategy (Fig. 6). We call for more researchers to shed light on the relationship between G-quadruplexes and coronaviruses. Only if we have a deeper understanding of coronaviruses can we better cope with the possible novel coronavirus pandemics in the future.

**Fig. 6.**
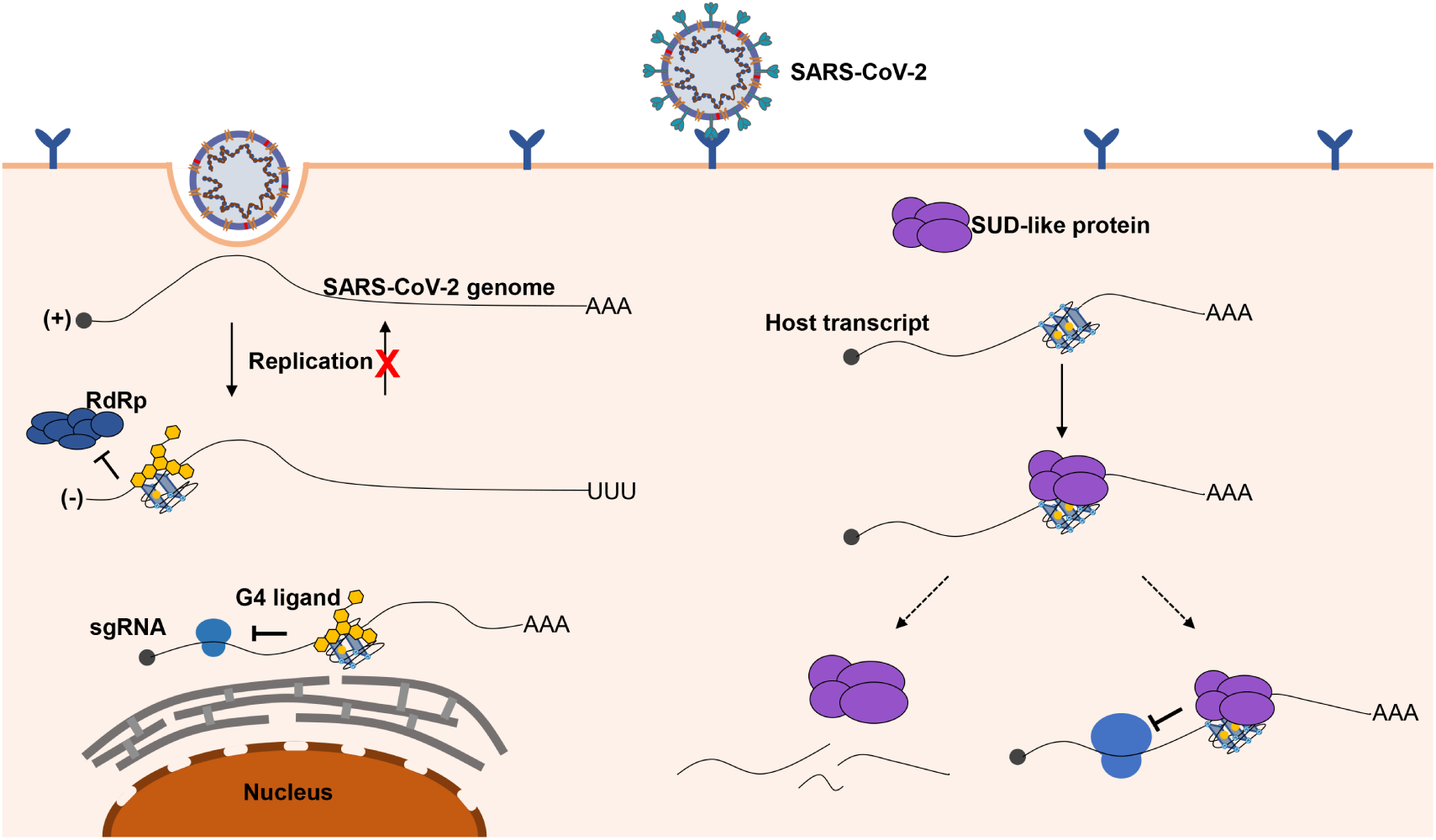
Possible role of G-quadruplexes in the antiviral mechanism and pathogenicity. Left part, G-quadruplexes can function as inhibition elements in the SARS-CoV-2 life cycle. Both the replication and translation could be affected by the G-quadruplexes structures. The stable G-quadruplexes in the 3’ end of the negative-sense strand may interfere with the activity of RdRp; hence, the replication of the negative-sense strands to the positive-sense strands is repressed, so that the SARS-CoV-2 genomes cannot be produced in large quantities. The G-quadruplex structures can suppress the translation process by impairing the elongating of ribosomes, which can hinder the production of proteins required for the virus. The G-quadruplex structures could be stabilized by the specific ligands to enhance the inhibitory effects, which is a promising antiviral strategy. Right part, a possible mechanism for SARS-CoV-2 to impede the expression of human genes. G-quadruplex structures, particularly with longer G-stretches, are the potential binding targets for SUD-like proteins. And the interaction of the SUD-like proteins with G-quadruplex structures possibly lead to the instability of host transcripts or obstructing the translation efficiency.

## Methods

### Data collection

We obtained a total of 77 full-length bat-associated beta coronaviruses from the DBatVir (http://www.mgc.ac.cn/DBatVir/) database[76]. We also downloaded the bat coronavirus RaTG13 genome from the NCBI virus database (https://www.ncbi.nlm.nih.gov/labs/virus/vssi/#/), which has shown a high sequence similarity to the SARS-CoV-2 reference genome in previous reports. We acquired the SARS-CoV-2 reference genome from the NCBI virus database under the accession number of NC_045512. In addition to those sequences, nine pangolin coronaviruses were derived from GISAID (https://www.gisaid.org/) database[77].

### Pairwise and multiple sequence alignment, phylogenetic and conservation analysis

The EMBOSS Needle software, which is based on the Needleman-Wunsch algorithm and, is a part of the EMBL-EBI web tools[78], was employed for the pairwise sequence alignment. Clustal Omega[79, 80] is a reliable and accurate multiple sequence alignment (MSA) tool that can be performed on large data sets. We choose this MSA tool for the alignment of viral genomes and the alignment of protein sequences under the default paraments. UGENE[81] is a powerful and user-friendly bioinformatics software, and we choose UGENE to visualize the pairwise and multiple sequence alignment results. We used the MEGA X software[82] to construct the Neighbor-Joining phylogenetic tree with 1,000 bootstrap replications. To depict the conservation state for each nucleotide site, the GERP++ software[83] was applied to calculate the “Rejected Substitutions” score column by column, which can reflect the constraints strength for each nucleotide sites.

### Potential G-quadruplex detection

Several open-source G-quadruplex detection software was used to search the PG4s both in the SARS-CoV-2 positive-sense and negative-sense strands. G4CatchAll[84], pqsfinder[85], and QGRS Mapper[86] were employed to predict the putative G-quadruplexes, respectively; Please see ref[87] for more information about the comparison of those tools mentioned above. The minimum G-tract length was set to two in the three software, while the max length of the predicted G-quadruplexes was limited to 30. Specifically, the minimum score of the predicted G-quadruplex was set to 10 when using pqsfinder. We utilized BEDTools[88] to sort the PG4s according to their coordinates. Apart from this, we adopted the cG/cC scoring system[54] proposed by Jean-Pierre Perreault et al. to delineate the sequence context influence on PG4s. The PG4s along with 15 nt upstream and downstream sequence contexts were used to calculate cG/cC score, and 2.05 was taken as the threshold for the preliminary inference of the G-quadruplex folding capability[54]. Using a customized python script, we implemented the cG/cC scoring system.

### Homo-dimer homology modeling and electrostatic potential calculation

The SARS-CoV-2 SUD_core_-like homo-dimer structure was modeled based on the template of the SARS-CoV SUD structure (PDB ID: 2W2G) through homology modeling. All the modeling process were performed in the Swiss Model[89] website (https://swissmodel.expasy.org/). The electrostatic potential was calculated and visualized in the PyMOL software by using the APBS (Adaptive Poisson-Boltzmann Solver) plugin.

### ΔG° z-score analysis

The ΔG° z-score for the SARS-CoV-2 genome was retrieved from RNAStructuromeDB (https://structurome.bb.iastate.edu/sars-cov-2). The ΔG° z-score is described as follows.

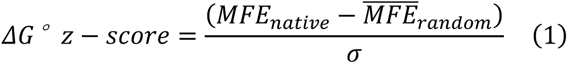

Where the *MFE*_*native*_ means the MFE (minimum free energy) ΔG° value predicted by the RNAfold software with a window of 120 nt and step of one nt. And the 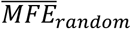 represents the MFE ΔG° value generated by the randomly shuffled sequence with the identical nucleotide composition. The *σ* is the standard deviation across all the MFE values.

To depict the ΔG° z-score for each nucleotide in the SARS-CoV-2 genome, we utilized the following formula.

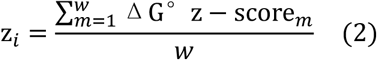

Where *z*_*i*_ is the average ΔG° z-score for nucleotide *i, w* denotes the total number of the sliding windows that covering the nucleotide *i*. ΔG° *z* − score_*m*_ indicates the ΔG° z-score for the m-th window. For example, when considering the nucleotide 1000 under the setting of 120 nt window length and one nt step, there are 120 sliding windows covering the nucleotide 1000. So, the *z*_200_, which means the average ΔG° z-score for nucleotide 200, is calculated as the sum of theΔG° z-score of 120 sliding windows divided by the total number of the sliding windows.

## Supplementary

**Fig. S1.**
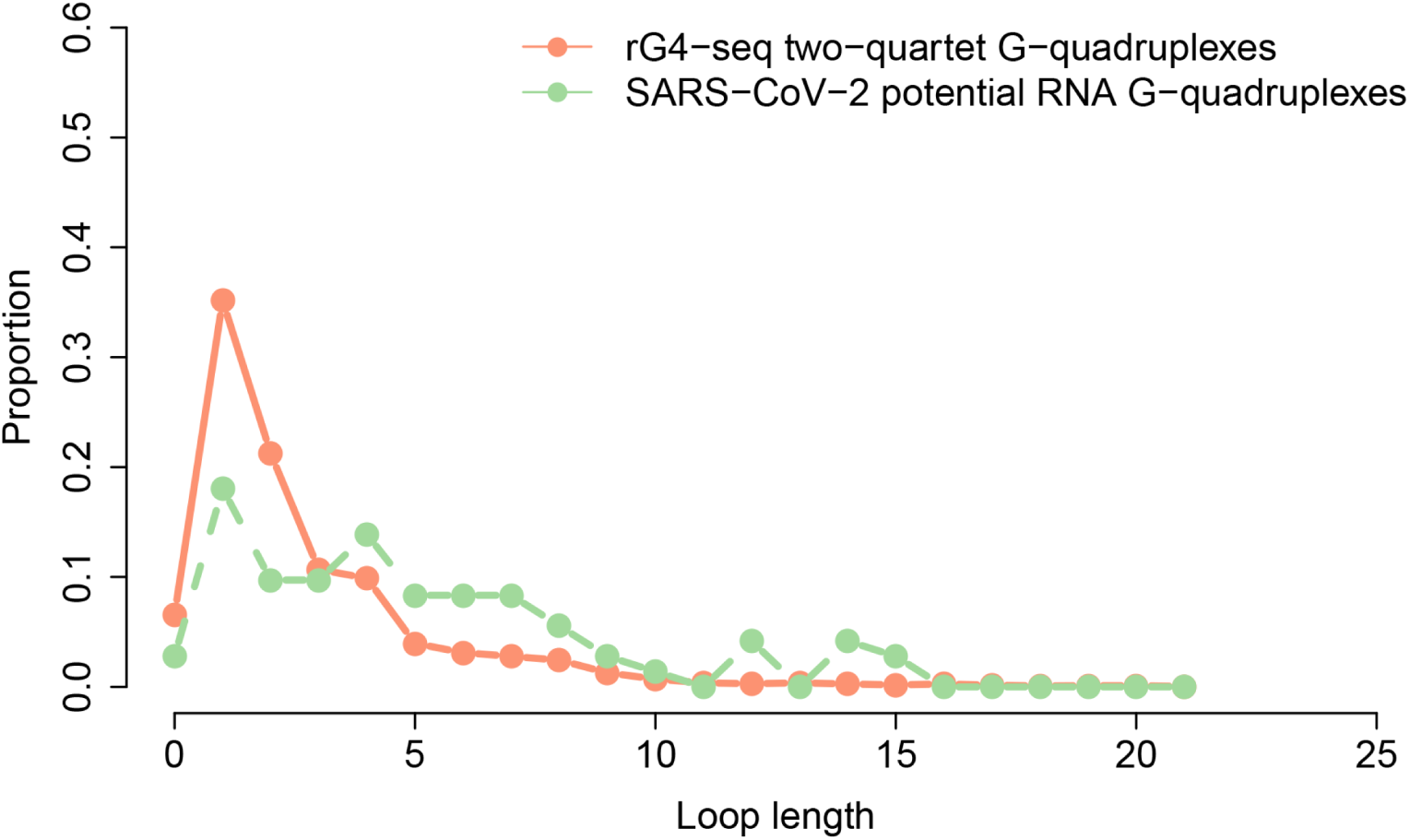
Proportion of loops with different lengths. The x-axis and y-axis show the loop length and loop proportion, respectively. The blue curve represents the loop in potential G-quadruplexes, while the red curve indicates the loop in two-quartet G-quadruplexes derived from rG4-seq.

**Fig. S2.**
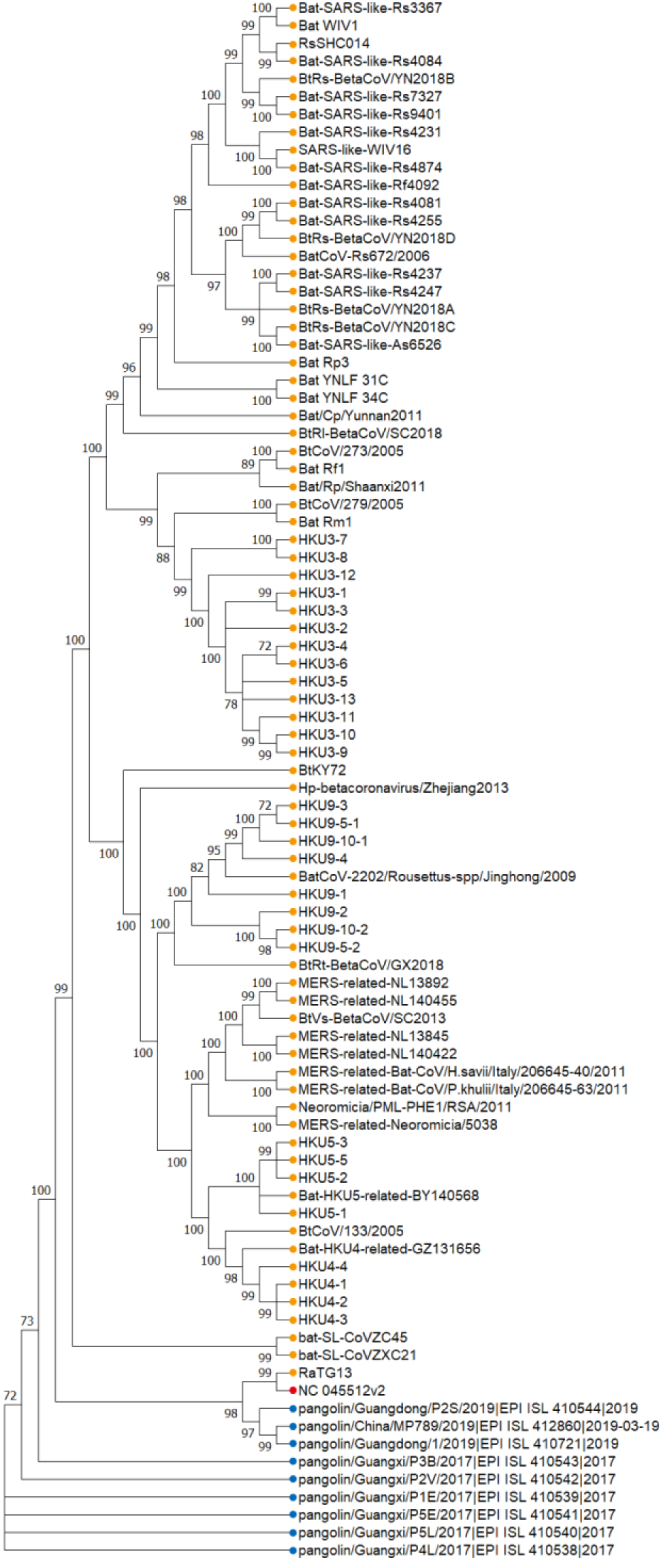
Phylogenetic tree of bat and pangolin related beta-coronavirus. The phylogenetic tree was constructed using the Neighbor-Joining method with 1,000 bootstrap replications. The bootstrap values lower than 70 were removed from the phylogenetic tree nodes. The SARS-CoV-2 reference sample is marked in the red dot, and the orange dots indicate the bat-related beta coronavirus samples, while the blue dots represent pangolin related beta-coronavirus samples.

**Fig. S3.**
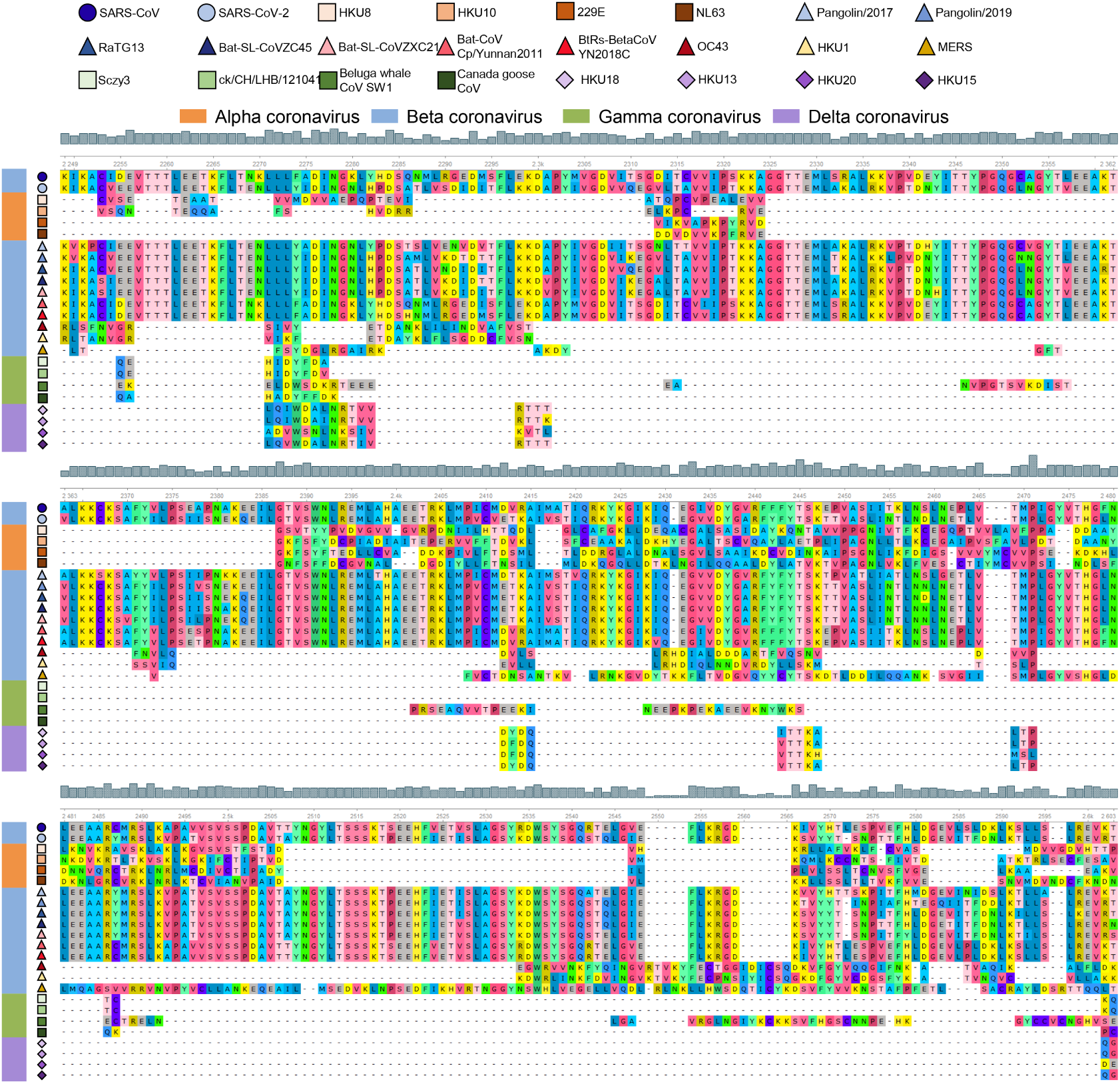
Alignment of several coronavirus amino acid sequences. The figure shows the sequence alignment corresponding to SARS SUD. The shapes in various colors mark different kinds of coronaviruses. The color bar represents different genera of coronaviruses (orange, alpha coronavirus; blue, beta coronavirus; green, gamma coronavirus; purple, delta coronavirus). The grey histogram shows the consensus of the alignment sites.

